## Supplementary material for "Receptor tyrosine kinases CAD96CA and FGFR1 function as the cell membrane receptors of insect juvenile hormone": Figure supplement, Supplementary files: Supplementary information-bioRxiv.pdf

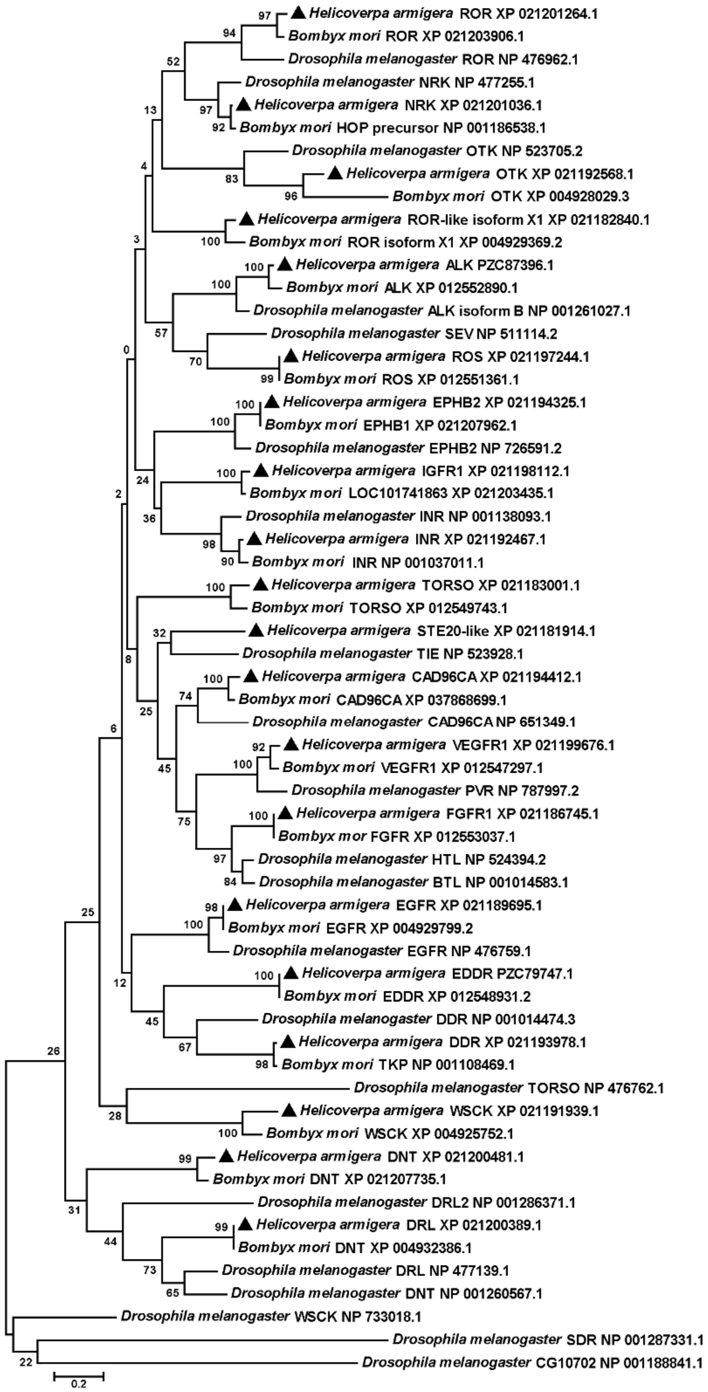


**Figure 1—figure supplement 1.** Phylogenetic tree analysis to identify RTKs of *H. armigera*. The phylogenetic tree was analyzed with MEGA 5.0, Corresponding amino acid sequences in RTKs of *H. armigera*, *B. mori*, and *D. melanogaster* obtained from NCBI. The tree shows clustering and the clades of various RTK in *H. armigera*, *B. mori*, and *D. melanogaster*. Black triangles represent RTKs in *H. armigera*. NRK was renamed based on the phylogenetic tree. The other RTKs were named based on the *H. armigera* genome.

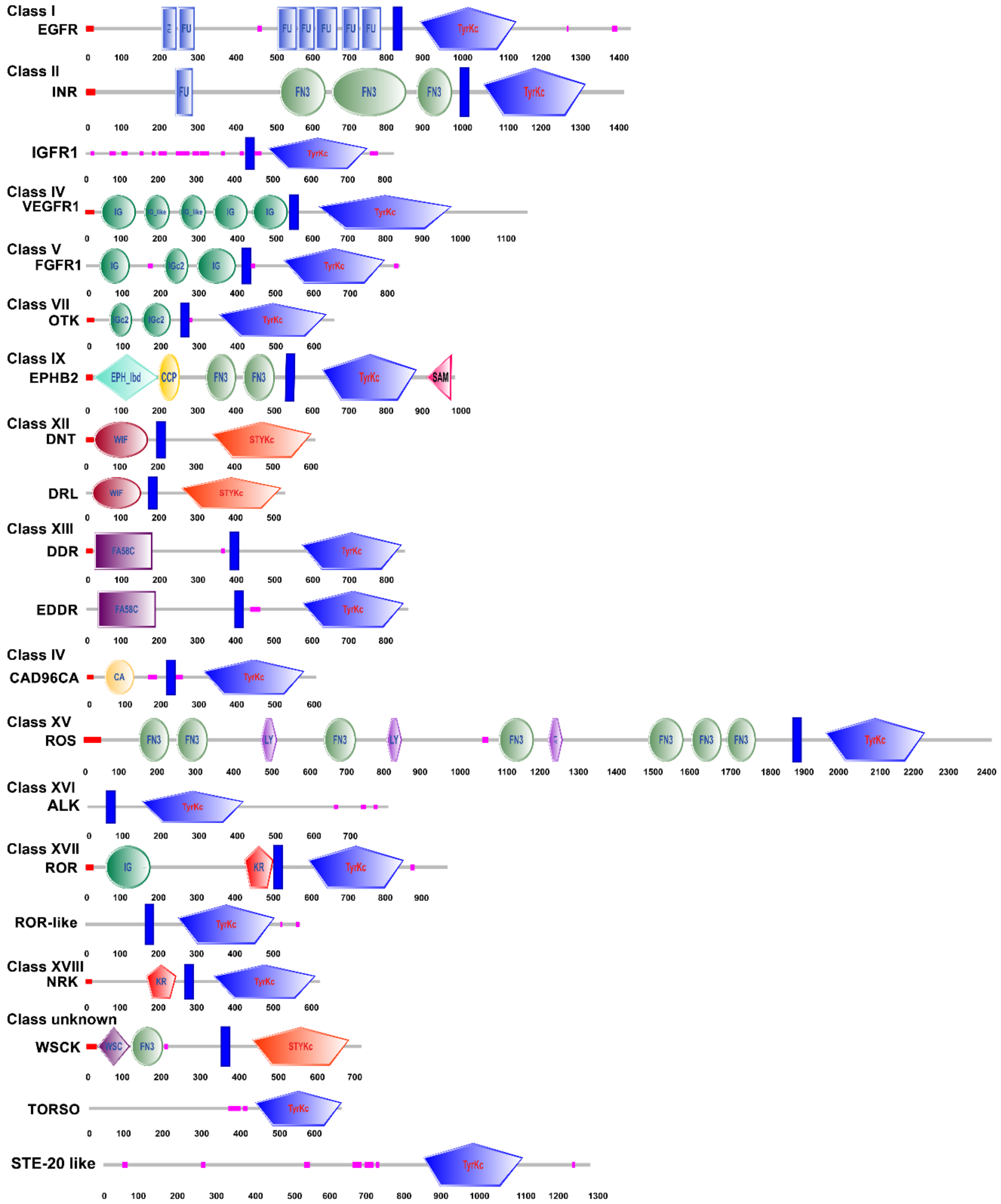


**Figure 1—figure supplement 2.** Structural characteristics of the RTK domains. The SMART tool was used to analyze the RTKs of *H. armigera.*

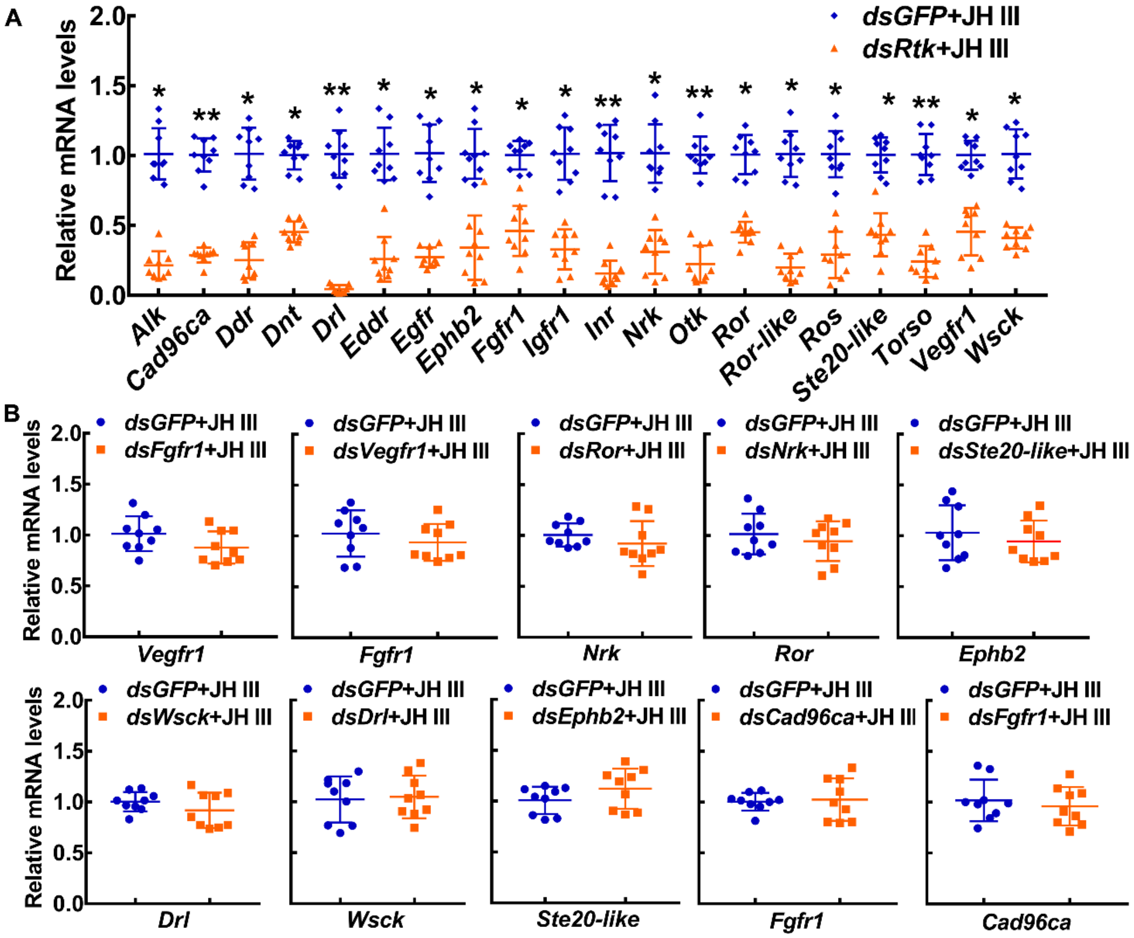


**Figure 1—figure supplement 3.** The interference efficiency of dsRNA and off–target detection. (**A**) The interference efficiency of dsRNA in HaEpi cells. (**B**) The qRT–PCR was performed to analyze the off–target genes. All of the relative mRNA levels were calculated via the $2^{-\Delta\Delta CT}$ method, and the bars indicate the mean ± SD according to three biological replicates and three technical replicates. Asterisks manifest significant differences by Student's *t* test (*$p$ < 0.05; **$p$ < 0.01).

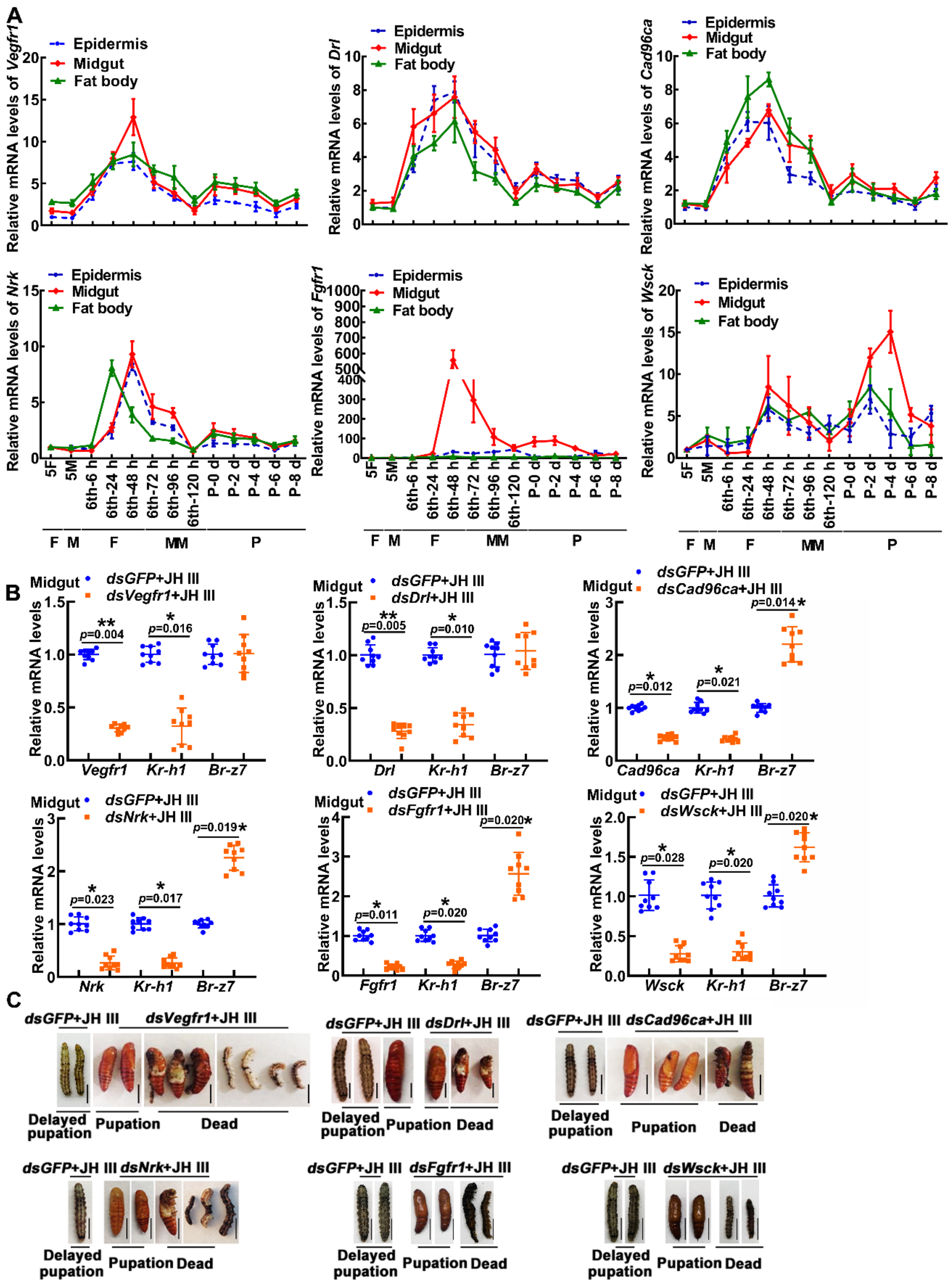


**Figure 1—figure supplement 4.** Expression profiles, interference efficiency and phenotype of 6 *Rtks* in larvae. (**A**) The expression profiles of *Vegfr1*, *Drl*, *Cad96ca*, *Nrk*, *Fgfr1*, and *Wsck* during

development. 5F: fifth instar feeding larvae; 5M: fifth instar molting larvae; 6th-6 h to 6th-120 h: sixth instar at 6 h to sixth instar 120 h larvae; P0 d to P8 d: pupal stage at 0-day to pupal stage at 8-day F: feeding stage; M: molting stage; MM: metamorphic molting stage; P: pupae. (**B**) qRT–PCR showed the interference efficiency of *Vegfr1*, *Drl*, *Cad96ca*, *Nrk*, *Fgfr1*, and *Wsck*, and the mRNA level of *Kr-h1* and *Br-z7*. The relative mRNA levels were calculated via the $2^{-\Delta\Delta CT}$ method, and the bars indicate the mean ± SD. Asterisks manifest significant differences by Student's *t* test (**p* < 0.05; ***p* < 0.01) based on three biological replicates, n = 3. (**C**) The phenotype after *Vegfr1*, *Drl*, *Cad96ca*, *Nrk*, *Fgfr1*, and *Wsck* knockdown. Scale bars: 1 cm.

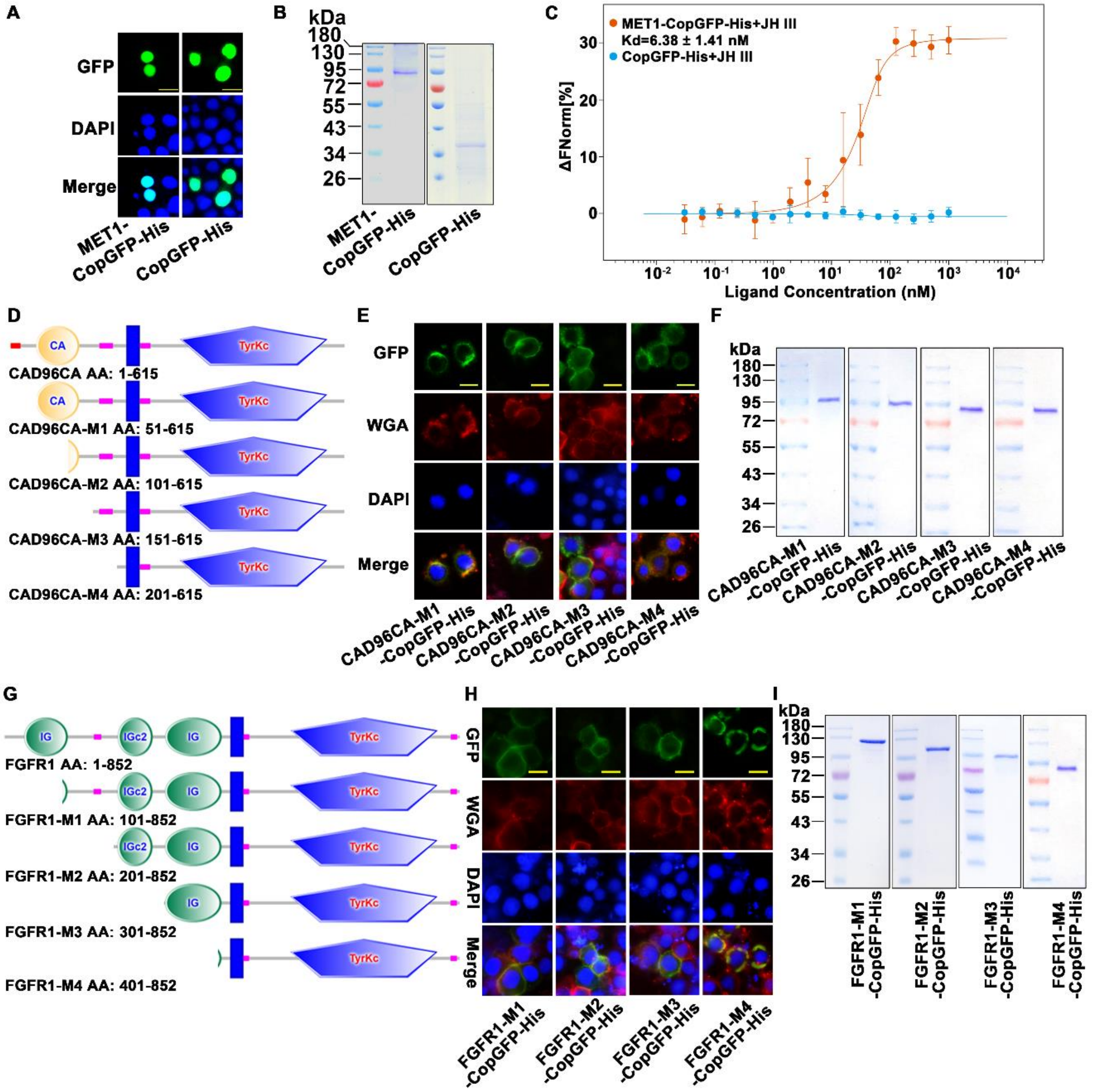


**Figure 3—figure supplement 1.** MET1 bound JH III, and CAD96CA and FGFR1 mutants. (**A**) The subcellular localization of overexpressed MET1-CopGFP-His and CopGFP-His in the cells. Green: green fluorescence from MET1-CopGFP-His and CopGFP-His. DAPI: nuclear staining. Merge: the pictures of different fluorescence-labelled cells were combined. The cells were observed with a fluorescence microscope. Scale bar=20 μm. (**B**) Coomassie brilliant blue staining of SDS–PAGE gel showing the purity of the separated MET1-CopGFP-His and CopGFP-His proteins. (**C**)

Saturation binding curves of MET1-CopGFP-His and CopGFP-His. Data are mean ± SE of three replicates. (**D**) The diagram of CAD96CA mutation. (**E**) Subcellular localization of the CAD96CA mutants. Green: green fluorescence of mutants fused with a green fluorescent protein. Red: the cell membrane stained with wheat germ lectin (WGA). Blue: nuclei stained with DAPI. Scale = 20 μm. (**F**) Coomassie brilliant blue staining of the SDS–PAGE gel showed the purity of the separated CAD96CA mutant proteins. (**G**) The diagram of FGFR1 mutation. (**H**) Subcellular localization of the FGFR1 mutants. Green: green fluorescence of mutants fused with a green fluorescent protein. Red: the cell membrane stained with wheat germ lectin (WGA). Blue: nuclei stained with DAPI. Scale = 20 μm. (**I**) Coomassie brilliant blue staining of the SDS–PAGE gel showed the purity of the separated FGFR1 mutant proteins.

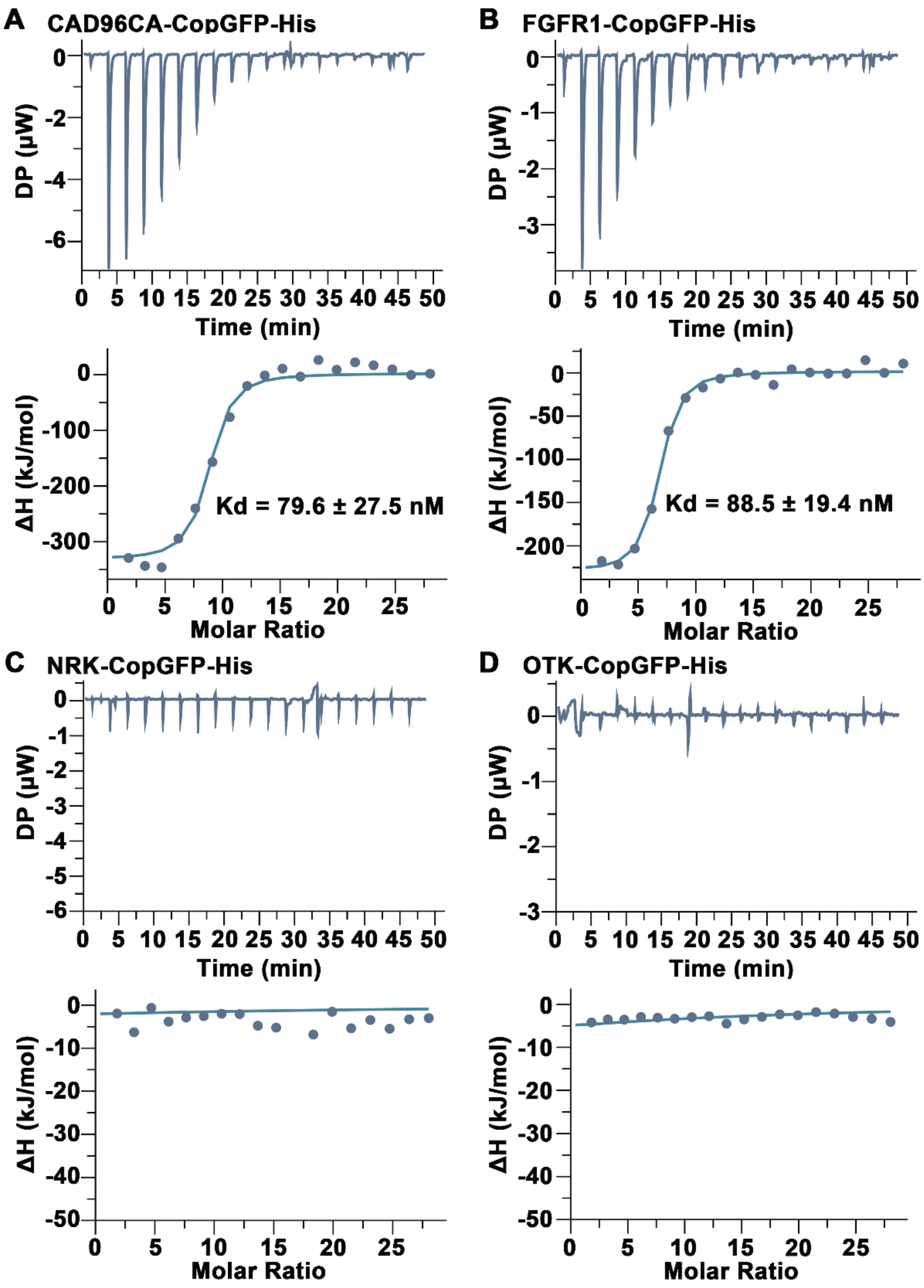


**Figure 3—figure supplement 2.** CAD96CA and FGFR1 bound JH III were analyzed using ITC. (**A**) Saturation binding curves of CAD96CA-CopGFP-His. (**B**) Saturation binding curves of FGFR1-CopGFP-His. (**C**) The binding curves of NRK-CopGFP-His. (**D**) The binding curves of OTK-CopGFP-His. The data were subtracted with that from the control test by the analysis software.

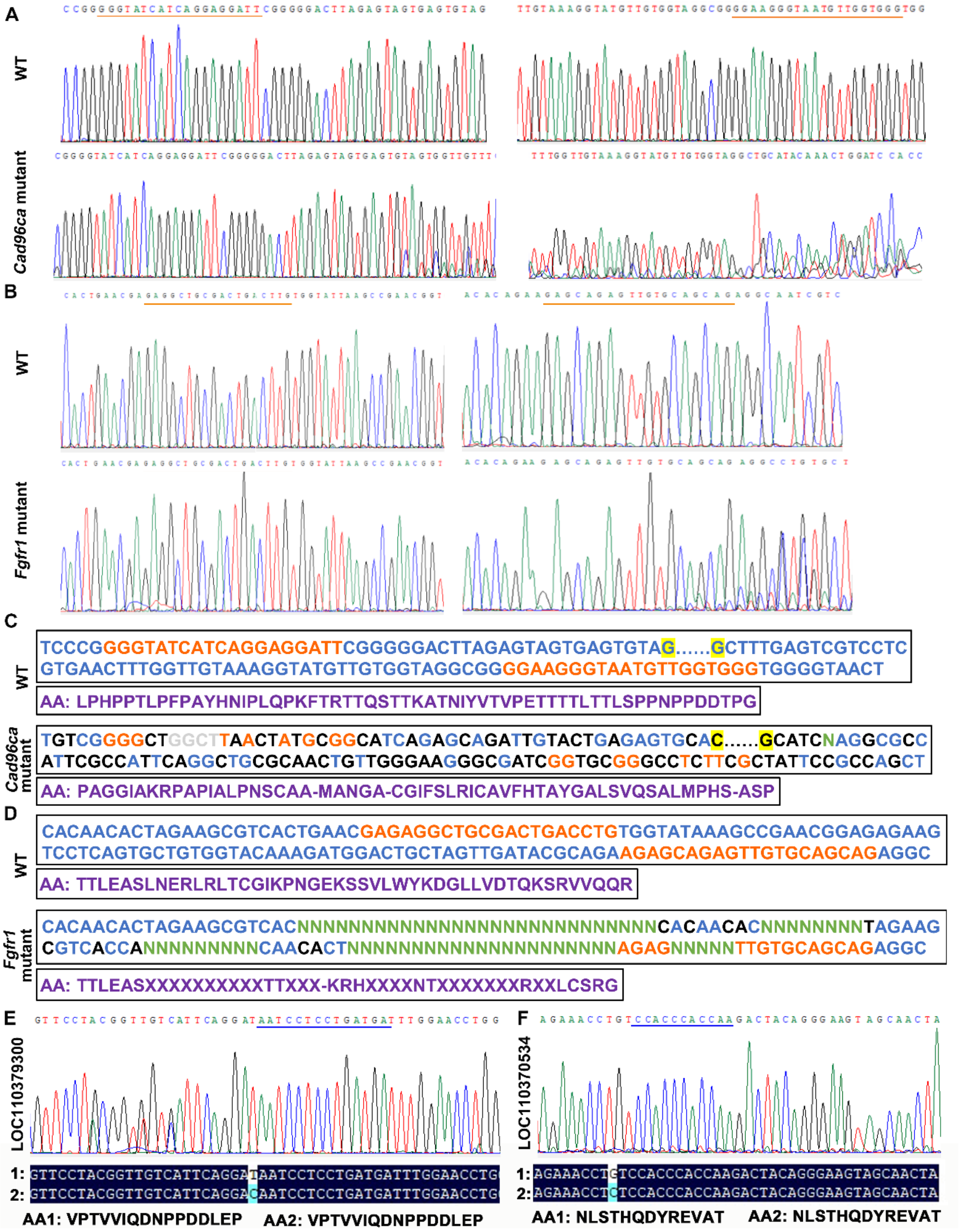


**Figure 4—figure supplement 1.** Targeted mutagenesis of *Cad96ca* and *Fgfr1* in *H. armigera*. (**A** and **B**) Mutations were detected by Sanger sequencing. Representative chromatograms of the PCR products of G0 *H. armigera* showing mutations induced by the CRISPR/Cas9 system. The gRNA target sequence was marked with an orange line. (**C** and **D**) Examples of G0 mutations identified by TA cloning and Sanger sequencing. The gRNA target sequence was marked in orange.

Nucleotide insertions were shown in grey; nucleotide deletions were shown in green (N); and substitutions were shown in black. The purple sequence represents the amino acid sequence. (**E** and **F**) Off–target genes detected by Sanger sequencing. The chromatograms of the PCR products of G0 *H. armigera* showed mutations. The gRNA target sequence was marked with a blue line. 1: mutant sequence, 2: normal sequence.

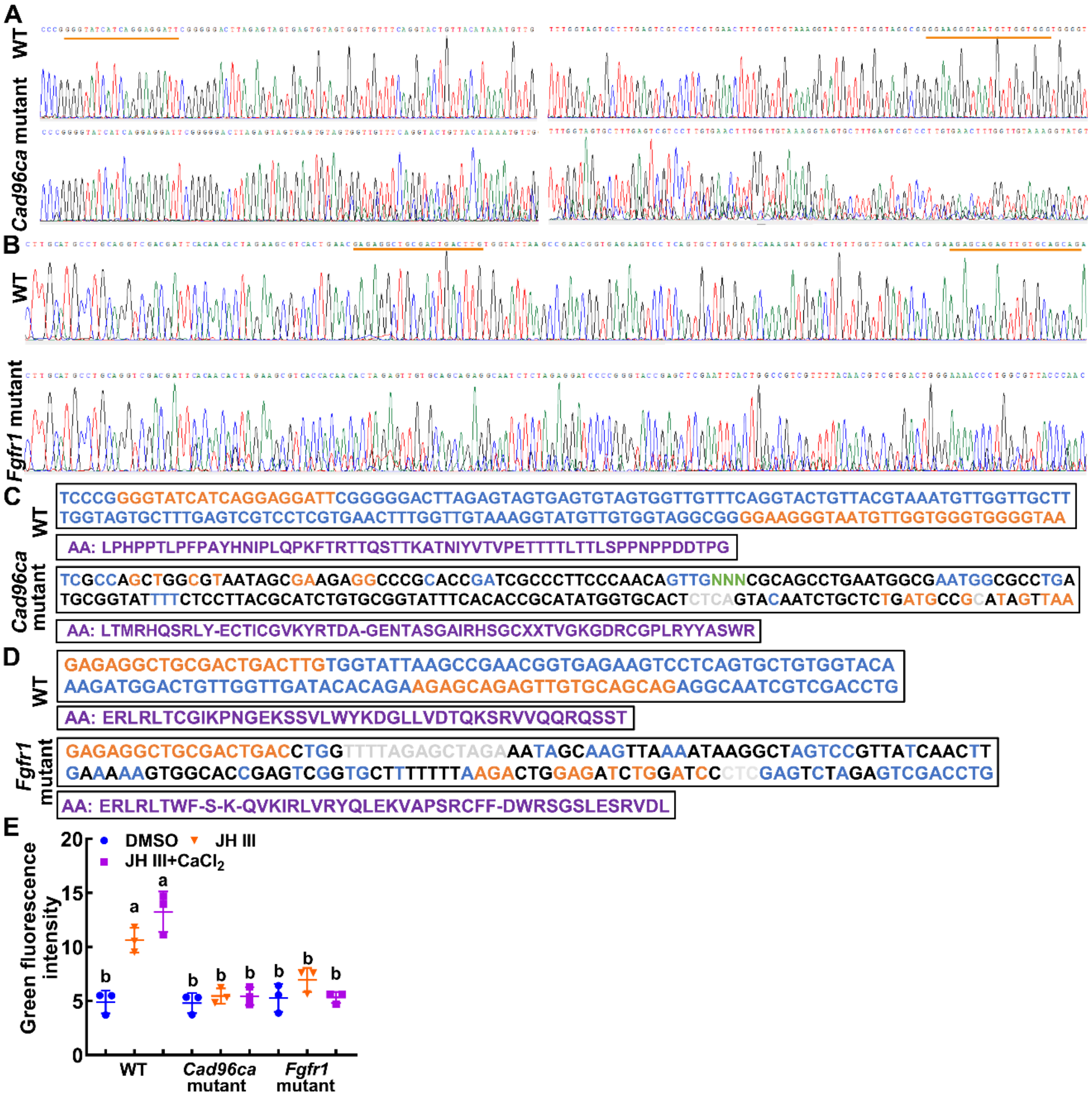


**Figure 4—figure supplement 2.** Targeted mutagenesis of *Cad96ca* and *Fgfr1* in HaEpi cells. (**A** and **B**) Mutations were detected by Sanger sequencing. Representative chromatograms of the PCR products of mutations. The gRNA target sequence was marked with an orange line. (**C** and **D**) Examples of mutations identified by TA cloning and Sanger sequencing. The gRNA target sequence was marked in orange. Nucleotide insertions were shown in grey, nucleotide deletions were shown in green (N), and substitutions were shown in black. The purple sequence represents the amino acid sequence. (**E**) Statistical analysis of the green fluorescence signal intensity by ImageJ software. The statistical analysis was performed using three independent replicates by ANOVA.

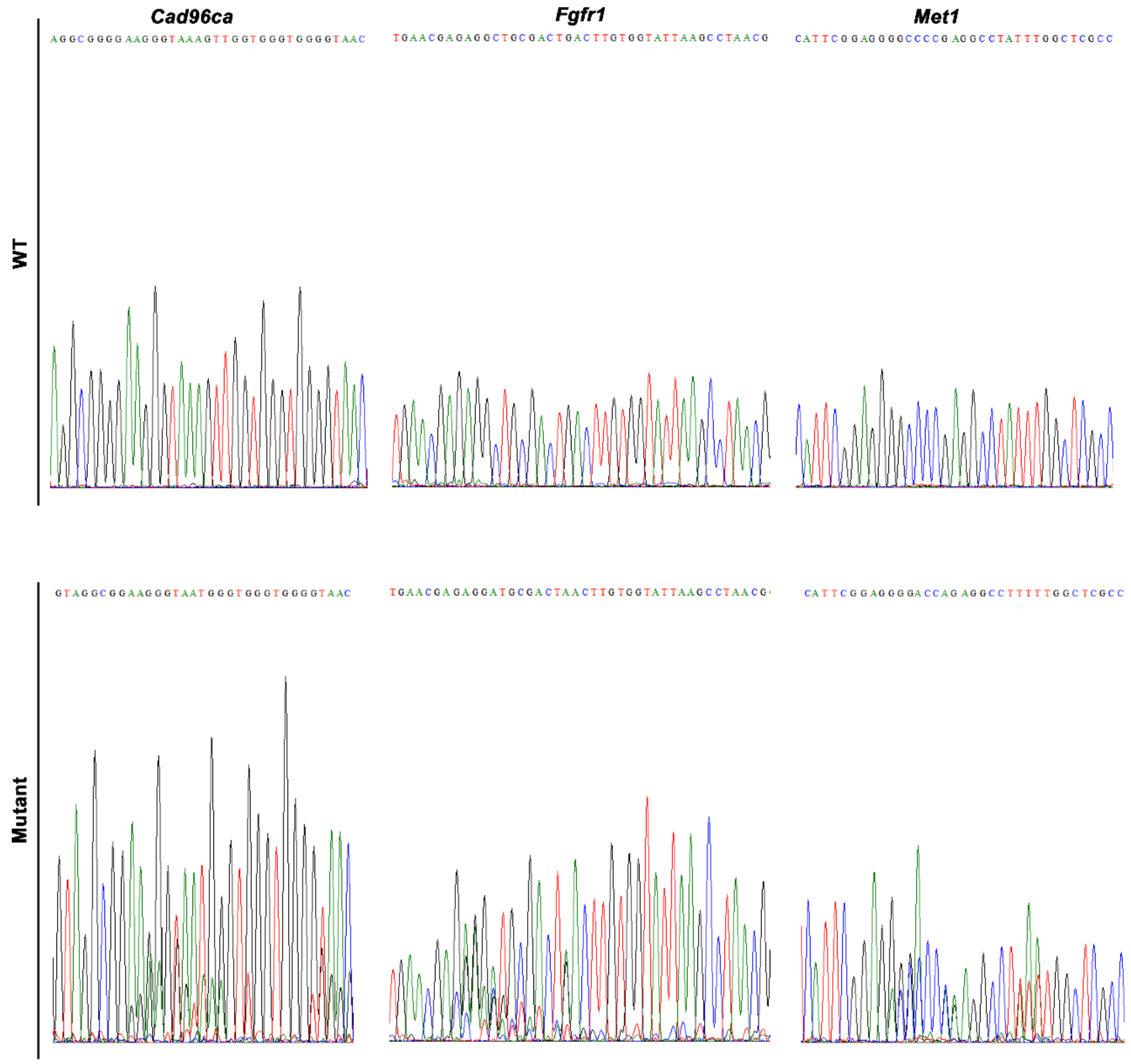


**Figure 5—figure supplement 1.** Targeted mutagenesis of *Cad96ca*, *Fgfr1*, and *Met1* in *H. armigera*. Mutations were detected by Sanger sequencing. Representative chromatograms of the PCR products of G0 *H. armigera* showing mutations induced by the CRISPR/Cas9 system.

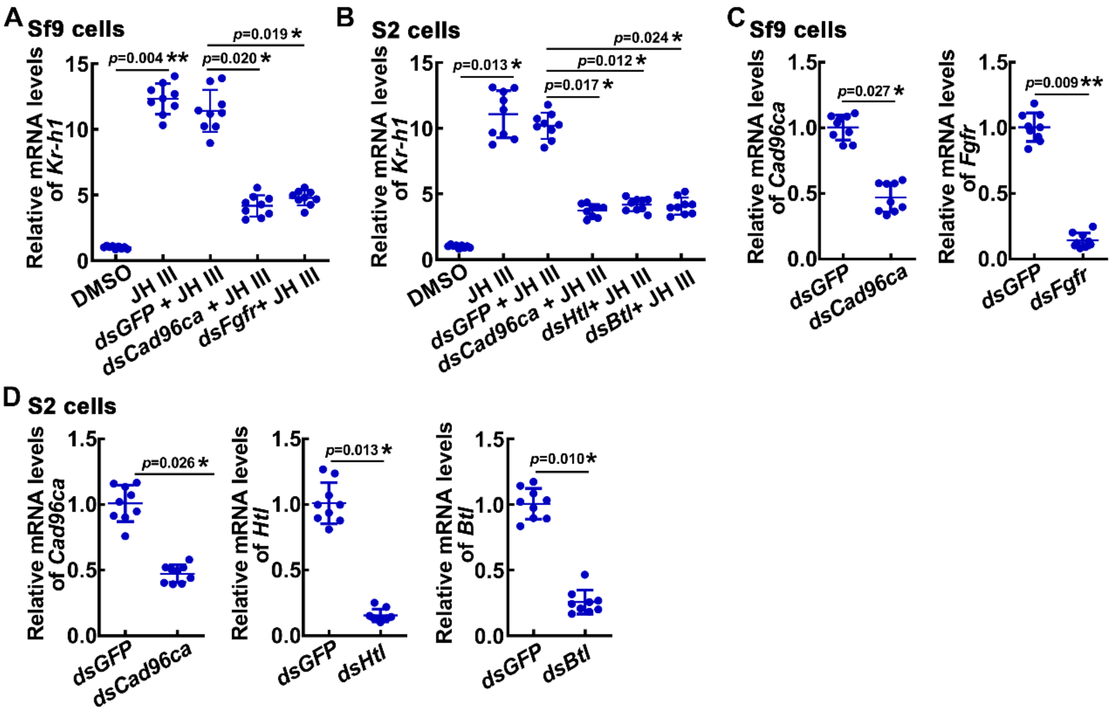


**Figure 6—figure supplement 1.** The gene expression was analyzed by qPCR. (**A**) The mRNA level of *Kr-h1* after genes knockdown in Sf9 cells. (**B**) The mRNA level of *Kr-h1* after genes knockdown in S2 cells. (**C**) Interference efficiency of *Cad96ca* and *Fgfr* in sf9 cell lines. (**D**) Interference efficiency of *Cad96ca*, *Htr* and *Btl* in S2 cell lines. The relative mRNA levels were calculated via the $2^{-\Delta\Delta CT}$ method, and the bars indicate the mean ± SD. Asterisks manifest significant differences by Student's *t* test (**p* < 0.05; ***p* < 0.01) based on three biological replicates, n = 3.

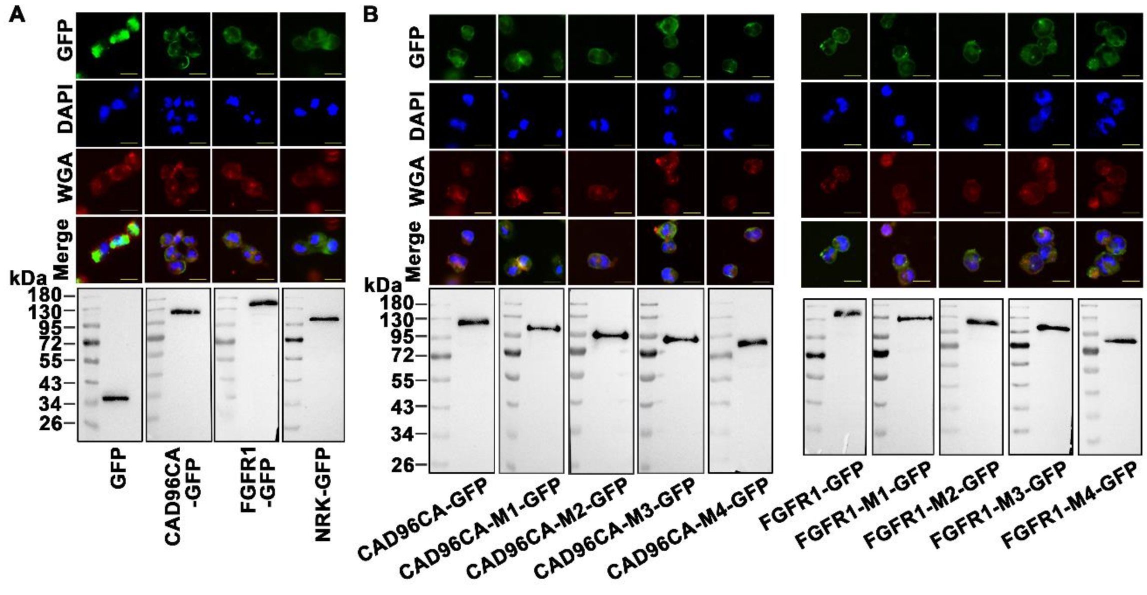

**Figure 6—figure supplement 2.** CAD96CA, FGFR1 and mutants overexpressed in HEK-293T cells. (**A**) Subcellular localization of overexpressed GFP, CAD96CA-GFP, FGFR1-GFP, and NRK-GFP. GFP: green fluorescence of RTKs fused with a green fluorescent protein. WGA: wheat germ agglutinin, a cell membrane label. DAPI: nuclear staining. Merge: the pictures of different fluorescence-labelled cells were combined. The cells were observed with a fluorescence microscope. Scale bar = 20 μm. Western blotting showed the expression of the target protein. (**B**) Subcellular localization of the CAD96CA and FGFR1 mutants. Green: green fluorescence of mutants fused with a green fluorescent protein. Red: the cell membrane stained with wheat germ lectin (WGA). Blue: nuclei stained with DAPI. Scale = 20 μm. The protein level of the mutant was detected by western blotting.

**Supplementary files**

Supplementary file 1. Names of RTKs identified in *H. armigera* genome.

| *Helicoverpa armigera* | | *Bombyx mori* | *Drosophila melanogaster* |
|---|---|---|---|
| Name | Symbol | Symbol | Symbol |
| ALK/Anaplastic lymphoma kinase | LOC110383585 | ALK | ALK |
| Cad96Ca/Cadherin 96Ca | LOC110379194 | Cad96Ca | Cad96Ca |
| Ddr/Discoidin domain receptor | LOC110378887 | TKP | Ddr |
| Dnt/Doughnut on | LOC110383864 | Dnt isoform X1 | Dnt |
| Drl/Derailed | LOC110383805 | Dnt | Drl |
| EDdr/Epithelial discoidin domain receptor | LOC110374488 | EDdr | |
| EGFR/Epidermal growth factor receptor | LOC110375773 | EGFR | Egfr |
| EphB2/Ephrin type-B receptor 2 | LOC110379128 | EphB1 | EphB2 |
| FGFR1/Fibroblast growth factor receptor homolog 1 | LOC110373728 | FGFR | Htl/DFR1/Dtk1 |
| IGFR1/Insulin-like growth factor 1 receptor | LOC110381988 | LOC101741863 | |
| InR/Insulin-like receptor | LOC110377777 | InR | InR |
| Nrk/Neurotropic receptor kinase | LOC110384207 | HOP | Nrk |
| Otk/Offtrack | LOC110377855 | Otk | Otk |
| Ror/Receptor tyrosine kinase orphan receptor | LOC110384348 | Ror | Ror |
| Ror-like isoform X1/ Receptor tyrosine kinase like orphan receptor | LOC110371076 | Ror isoform X1 | Ror |
| ROS/Proto-oncogene tyrosine-protein kinase | LOC110381275 | ROS | Sev |
| STE20-like/serine/ | LOC110370444 | | |

| | | | |
|---|---|---|---|
| threonine-protein kinase STE20-like | | | |
| Torso/tyrosine-protein kinase receptor torso like | LOC110371197 | Torso | Torso |
| VEGFR1/Vascular endothelial growth factor receptor 1 | LOC110383235 | VEGFR1 | Pvr |
| Wsck/Cell wall integrity and stress response component kinase | LOC110377380 | Wsck | Wsck |

Supplementary file 2. Oligonucleotide sequences of PCR primers.

| Primer name | 5´⟶3´ nucleotide sequence |
|---|---|
| **qRT-PCR** | |
| Kr-h1-RTF | atgtttacgagatttcggttac |
| Kr-h1-RTR | atgtgggcttccatttgtttt |
| Jhi-1-RTF | accacatcttcatcacaacca |
| Jhi-1-RTR | tacaactcatccaagccctca |
| Jhi-26-RTF | gcggatacgaaccacat |
| Jhi-26-RTR | ggctccactgacacgat |
| Vg-RTF | gtcaatgaggatgaacaggga |
| Vg-RTR | gttggcgttagacacgagagg |
| Torso-RTF | cgggcagataagcacaactc |
| Torso-RTR | gaggaaaggctcgtttgatg |
| Otk-RTF | gtgcgtgattcgttcgtt |
| Otk-RTR | ccttctactcgacttgtggg |
| Ddr-RTF | gtgtccgaggtcgcaaat |
| Ddr-RTR | cgataacatacgcctctgc |
| Wsck-RTF | gattggagtggtggcagtt |
| Wsck-RTR | tgtggttgccaagggtat |
| Egfr-RTF | gactatctgatgccctcaccgc |
| Egfr-RTR | aaccgcaaatcctttattccct |
| Ste20-like-RTF | ctcgccacgctactccaca |
| Ste20-like -RTR | tcatactccgccgacagg |
| Vegfr1-RTF | ttaggttgaaagattacccacg |
| Vegfr1-RTR | atctccagtacgctcgtgtc |
| Ror-like-RTF | tcacgcacgaatcagacg |
| Ror-like-RTR | tggcggcacaagcacta |
| Fgfr1-RTF | gtggcaacggcgtgtctt |
| Fgfr1-RTR | aactctgctcttctgcgtatca |
| Ros-RTF | tcccgctcgtgagtatga |
| Ros-RTR | tgattgagtgttccgtgctat |
| Igfr1-RTF | tgctgctgtgcctgctggtg |
| Igfr1-RTR | cggtgccgagtttccgatta |
| Inr-RTF | tcttggtacaccgtgaacatc |
| Inr-RTR | actacgaagccgttggggttctgag |
| Dnt-RTF | cgagaaactaaggctgaaggtg |
| Dnt-RTR | gccagaggtgatgctccaag |
| Drl-RTF | agatgcgagggagcaagaagt |
| Drl-RTR | gctaacacccaggaccgacag |
| Cad96ca-RTF | ttcaacctacccgccatca |
| Cad96ca-RTR | tctccaacccataagtcacag |
| Alk-RTF | aagaaggcggtgatagacgatt |
| Alk-RTR | tgactgttggacgaggaggac |
| Nrk-RTF | ggactacagccaagtaaccac |

| | |
|---|---|
| Nrk-RTR | gaggtcttgtatgctgatgagggta |
| Ror-RTF | acacgccgcaaaggagac |
| Ror-RTR | ccgttggaagaggagcag |
| Ephb2-RTF | cagtgctggagacaaccttcg |
| Ephb2-RTR | tcggctgtttcttatcacattca |
| Eddr-RTF | atgcgacctgtcaccttccttg |
| Eddr-RTR | tgccgctttcacttcgttatgg |
| **RNAi** | |
| Fgfr1-RNAiF | gcgtaatacgactcactatagggagcgtcactgaacgagag |
| Fgfr1-RNAiR | gcgtaatacgactcactatagggaaacgtggagggaaatat |
| Vegfr1-RNAiF | gcgtaatacgactcactatagggttgcctcacttcagcc |
| Vegfr1-RNAiR | gcgtaatacgactcactatagggtttcgcactttccacg |
| Wsck-RNAiF | gcgtaatacgactcactataggg tttctgtgggaatgcg |
| Wsck-RNAiR | gcgtaatacgactcactatagggggctggggtctggagt |
| Drl-RNAiF | gcgtaatacgactcactataggggagtggacttgccttgtacg |
| Drl-RNAiR | gcgtaatacgactcactatagggtcagctctgctatcctttgt |
| Cad96ca-RNAiF | gcgtaatacgactcactataggggtctacgccacagtctccga |
| Cad96ca-RNAiR | gcgtaatacgactcactatagggcgtctttcttgctatccttc |
| Ror-RNAiF | gcgtaatacgactcactatagggggcgtgtatttattgttt |
| Ror-RNAiR | gcgtaatacgactcactatagggggtgccattagtcttatc |
| Ephb2-RNAiF | gcgtaatacgactcactatagggatacccactggctcctgt |
| Ephb2-RNAiR | gcgtaatacgactcactatagggcattctcggcgtaaactt |
| Nrk-RNAiF | gcgtaatacgactcactatagggttatcgtgcttcttctta |
| Nrk-RNAiR | gcgtaatacgactcactatagggatgttgtggttacttggc |
| Ste20-like-RNAiF | gcgtaatacgactcactataggggcagaaaagacctacacagc |
| Ste20-like-RNAiR | gcgtaatacgactcactatagggcaggcaagtaacgtcacaac |
| **Overexpression** | |
| Nrk-oveF | gattctagagctagcgaattcgccaccatggacattcactttaa |
| Nrk-oveR | tcgtcgctctccatagcggccgcttcaggatgagttctttccaatatca |
| Otk-oveF | gattctagagctagcgaattcgccaccatggtgatgtgcgtgattcgttcgttc |
| Otk-oveR | tcgtcgctctccatagcggccgcctcttcgactttctcctgagatttc |
| Cad96ca-oveF | gattctagagctagcgaattcgccaccatggtgatgtttctgacaagc |
| Cad96ca-oveR | tcgtcgctctccatagcggccgctagtttttctccatccaagtgctg |
| Fgfr1-oveF | gattctagagctagcgaattcgccaccatgaatctcgccg |
| Fgfr1-oveR | tcgtcgctctccatagcggccgctttgatgaaaggaaagtcactgtca |
| **Mutant** | |
| Cad96ca-M1-F | gattctagagctagcgaattcgccaccatggtgagggtgtaccgtgaag |
| Cad96ca-M1-R | tcgtcgctctccatagcggccgctagtttttctccatccaagtgctg |
| Cad96ca-M2-F | gattctagagctagcgaattcgccaccatggtgtgggtgacagcatacg |
| Cad96ca-M2-R | tcgtcgctctccatagcggccgctagtttttctccatccaagtgctg |
| Cad96ca-M3-F | gattctagagctagcgaattcgccaccatggtgaggacgactcaaagca |
| Cad96ca-M3-R | tcgtcgctctccatagcggccgctagtttttctccatccaagtgctg |
| Cad96ca-M4-F | gattctagagctagcgaattcgccaccatggtgacagaagctcctaata |

| | |
|---|---|
| Cad96ca-M4-R | tcgtcgctctccatagcggccgctagtttttctccatccaagtgctg |
| Fgfr1-M1-F | gattctagagctagcgaattcgccacctgtaagactgataat |
| Fgfr1-M1-R | tcgtcgctctccatagcggccgctttgatgaaaggaaagtcactgtca |
| Fgfr1-M2-F | gattctagagctagcgaattcgccacccaccctacaaaacttt |
| Fgfr1-M2-R | tcgtcgctctccatagcggccgctttgatgaaaggaaagtcactgtca |
| Fgfr1-M3-F | gattctagagctagcgaattcgccaccgctgaaaacttgaccg |
| Fgfr1-M3-R | tcgtcgctctccatagcggccgctttgatgaaaggaaagtcactgtca |
| Fgfr1-M4-F | gattctagagctagcgaattcgccaccggatacttgactgtat |
| Fgfr1-M4-R | tcgtcgctctccatagcggccgctttgatgaaaggaaagtcactgtca |
| Crispr-Cas9 mutant | |
| Universal primer | aaaagcaccgactcggtgccactttttcaagttgataacggactagccttattttaacttgctattt ctagctctaaaac |
| Cad96ca-gRNA1 | taatacgactcactataggaagggtaatgttggtggggttttagagctagaa |
| Cad96ca-gRNA2 | taatacgactcactatagggtatcatcaggaggattgttttagagctagaa |
| Cad96ca-gRNAF1 | aagtggaagggtaatgttggtggggt |
| Cad96ca-gRNAR1 | taaaaccccaccaacattacccttcc |
| Cad96ca-gRNAF2 | aagtgggtatcatcaggaggattgt |
| Cad96ca-gRNAR2 | taaaacaatcctcctgatgataccc |
| Cad96ca-testF | gacagaagtctacgccaca |
| Cad96ca-testR | gcatacaaacaggatcaca |
| Fgfr1-gRNA1 | taatacgactcactatagggaggctgcgactgacctggttttagagctagaa |
| Fgfr1-gRNA2 | taatacgactcactataggagcagagttgtgcagcaggttttagagctagaa |
| Fgfr1-gRNAF1 | aagtgagaggctgcgactgacctggt |
| Fgfr1-gRNAR1 | taaaaccaggtcagtcgcagcctctc |
| Fgfr1-gRNAF2 | aagtagagcagagttgtgcagcaggt |
| Fgfr1-gRNAR2 | taaaacctgctgcacaactctgctct |
| Fgfr1-testF | acccaataaacaacctca |
| Fgfr1-testR | ctggtccttctactatacttac |
| gRNAwf-F | tgattacgaattcccgggaggttatgtagtacacattg |
| gRNAwf-R | gtgttttacgcgcccgggaaaaaaagcaccgactcggt |
| Met1-gRNA | taatacgactcactatagaggggccccgaggcctattgttttagagctagaa |
| Met1-testF | atgacatcttcaggcggag |
| Met1-testR | ctatatcggacaacaaaga |
| **HEK-239T** | |
| Overexpression | |
| Cad96ca-W-F | ctcgagaccatggtggaattcatgtttctgacaagcgtctggg |
| Cad96ca-W-R | ctcgcccttgctcatggtacctagtttttctccatccaagtgctg |
| Cad96ca-M1-F | ctcgagaccatggtggaattcagggtgtaccgtgaaggcagt |
| Cad96ca-M1-R | ctcgcccttgctcatggtacctagtttttctccatccaagtgctg |
| Cad96ca-M2-F | ctcgagaccatggtggaattctgggtgacagcatacgacggc |
| Cad96ca-M2-R | ctcgcccttgctcatggtacctagtttttctccatccaagtgctg |
| Cad96ca-M3-F | ctcgagaccatggtggaattcaggacgactcaaagcactacc |
| Cad96ca-M3-R | ctcgcccttgctcatggtacctagtttttctccatccaagtgctg |
| Cad96ca-M4-F | ctcgagaccatggtggaattcacagaagctcctaataagaat |
| Cad96ca-M4-R | ctcgcccttgctcatggtacctagtttttctccatccaagtgctg |
| Fgfr1-W-F | ctcgagaccatggtggaattcatgaatctcgccgccattg |
| Fgfr1-W-R | ctcgcccttgctcatggtacctttgatgaaaggaaagtcactgtca |
| Fgfr1-M1-F | ctcgagaccatggtggaattctgtaagactgataatgataatg |
| Fgfr1-M1-R | ctcgcccttgctcatggtacctttgatgaaaggaaagtcactgtca |
| Fgfr1-M2-F | ctcgagaccatggtggaattccaccctacaaaactttacaaaat |

| | |
|---|---|
| Fgfr1-M2-R | ctcgcccttgctcatggtacctttgatgaaaggaaagtcactgtca |
| Fgfr1-M3-F | ctcgagaccatggtggaattcgctgaaaacttgaccgttgtag |
| Fgfr1-M3-R | ctcgcccttgctcatggtacctttgatgaaaggaaagtcactgtca |
| Fgfr1-M4-F | ctcgagaccatggtggaattcggatacttgactgtattggaat |
| Fgfr1-M4-R | ctcgcccttgctcatggtacctttgatgaaaggaaagtcactgtca |
| ***S. frugiperda*** | |
| **qRT-PCR** | |
| Cad96ca-RTF | tgaacttgaacttccctgcct |
| Cad96ca-RTR | tcaccacgagaactcctatgc |
| Fgfr-RTF | tggctccagagtcgctttat |
| Fgfr-RTR | ccaggtcttccaccagttca |
| Kr-h1-RTF | tctgtcttcgtgatttcggtta |
| Kr-h1-RTR | gcctccatttgtttttttgtgt |
| **RNAi** | |
| Cad96ca-RNAiF | gcgtaatacgactcactataggtgcctaccccagtttaccta |
| Cad96ca-RNAiR | gcgtaatacgactcactataggccgttgtcccaagactcata |
| Fgfr-RNAiF | gcgtaatacgactcactataggcgaaggaggatgaagggtg |
| Fgfr-RNAiR | gcgtaatacgactcactataggcgactggcattagcaaagc |
| ***D. melanogaster*** | |
| **qRT-PCR** | |
| Cad96ca-RTF | tcaaggaaagtgccaccgaagt |
| Cad96ca-RTR | gcagcccaacaaatgaacaaca |
| Htl-RTF | tcaaggaaagtgccaccgaagt |
| Htl-RTR | gcagcccaacaaatgaacaaca |
| Btl-RTF | tagtggctttgttccttttgg |
| Btl-RTR | acggttgttctctgcttctcg |
| Kr-h1-RTF | cctaccactgtgacatctgctt |
| Kr-h1-RTR | tccatctccttcttgttcttga |
| **RNAi** | |
| Cad96ca-RNAiF | gcgtaatacgactcactatagggggagagcagagtggagaa |
| Cad96ca-RNAiR | gcgtaatacgactcactataggcagggaacatagcgattgg |
| Htl-RNAiF | gcgtaatacgactcactatagggggagagcagagtggagaa |
| Htl-RNAiR | gcgtaatacgactcactataggcagggaacatagcgattgg |
| Btl-RNAiF | gcgtaatacgactcactataggggctacccttttgcgactt |
| Btl-RNAiR | gcgtaatacgactcactataggcaccgtgtgctttctgctt |
